# Unveiling the genetic blueprint of a desert scorpion: A chromosome-level genome of *Hadrurus arizonensis* provides the first reference for Parvorder Iurida

**DOI:** 10.1101/2024.03.22.586284

**Authors:** Meridia Jane Bryant, Asher M. Coello, Adam M. Glendening, Samuel A. Hilliman, Carolina Fernanda Jara, Samuel S. Pring, Aviel Rodriguez Rivera, Jennifer Santiago Membreño, Lisa Nigro, Nicole Pauloski, Matthew R. Graham, Teisha King, Elizabeth L. Jockusch, Rachel J. O’Neill, Jill L. Wegrzyn, Carlos E. Santibáñez-López, Cynthia N. Webster

## Abstract

Over 400 million years old, scorpions represent an ancient group of arachnids and one of the first animals to adapt to life on land. Presently, the lack of available genomes within scorpions hinders research on their evolution. This study leverages ultra-long nanopore sequencing and Pore-C to generate the first chromosome level assembly and annotation for the desert hairy scorpion, *Hadrurus arizonensis*. The assembled genome is 2.23 Gb in size with an N50 of 280 Mb. Pore-C scaffolding re-oriented 99.6% of bases into nine chromosomes and BUSCO identified 998 (98.6%) complete arthropod single copy orthologs. Repetitive elements represent 54.69% of the assembled bases, including 872,874 (29.39%) LINE elements. A total of 18,996 protein-coding genes and 75,256 transcripts were predicted, and extracted protein sequences yielded a BUSCO score of 97.2%. This is the first genome assembled and annotated within the family Hadruridae, representing a crucial resource for closing gaps in genomic knowledge of scorpions, resolving arachnid phylogeny, and advancing studies in comparative and functional genomics.

**Significance:** Genomic resources for the study of arachnids are limited. To date, only four scorpion genomes have been published; none of these are chromosome-level assemblies, and all four belong to a single family, Buthidae. In this study, we assembled the first chromosome-level, annotated genome for a non-buthid species (*Hadrurus arizonensis*). This high quality reference will provide a critical resource for comparative and functional genomics and contribute to the understanding of arachnid evolution.

## Introduction

Arachnids emerged on land over 400 million years ago (Coddington & Colwell 2001) and are a diverse taxonomic group containing over 100,000 known species (Proctor *et al*. 2015). They inhabit a range of habitats, including terrestrial and aquatic environments (Kuntner 2022). Despite this diversity, genomic research within Arachnida has primarily focused on Araneae (spiders) (Sanggaard *et al*. 2014) and Parasitiformes (ticks) (Pagel Van Zee *et al*. 2007).

The order Scorpiones comprises 23 families and over 2,800 species in two main parvorders: Buthida and Iurida (Santibáñez-López *et al*. 2022, 2023). Most are nocturnal, solitary predators and the majority of species fluoresce under UV light. Being Arachnopulmonates, they possess book lungs for respiration; however, unlike other arachnids, their body is uniquely segmented into a prosoma, mesosoma, and metasoma (Howard *et al*. 2019). They have a pair of chelate pedipalps for defense and prey acquisition (prosoma), ventral sensory organs called pectines (mesosoma), and a segmented tail that ends in a telson and stinger (metasoma) to deliver venom. Scorpion venom is often the subject of research for human health and biotechnological applications (Kerkis *et al*. 2017). Despite this, significant gaps persist in regards to the diversity and evolution of scorpions (Kuntner 2022).

The higher-level relationships among scorpions have been controversial, especially with regard to monophyly and trait evolution (Lozano-Fernandez *et al*. 2019; Santibáñez-López *et al*. 2020; Ballesteros *et al*. 2022). The contribution of more high-quality genomes from underrepresented families, apart from the four buthid genomes currently available, can improve our understanding of relationships among scorpions and the evolution of unique traits (Long *et al*. 2003; Rittschof & Robinson 2016).

This study presents the first chromosome-scale genome assembly for the desert hairy scorpion, *Hadrurus arizonensis*. This species is an iconic inhabitant of the Mojave and Sonoran deserts of North America, and one of the largest scorpions found on the continent (Figure 1A). It also provides the first genome assembly for any species of the parvorder Iurida (Figure 1B). The sequenced genome of *H. arizonensis*, known to be significantly impacted by historical climate shifts (Graham *et al*. 2013), also offers a valuable foundation for future studies of adaptation in extreme environments.

**Figure 1.**
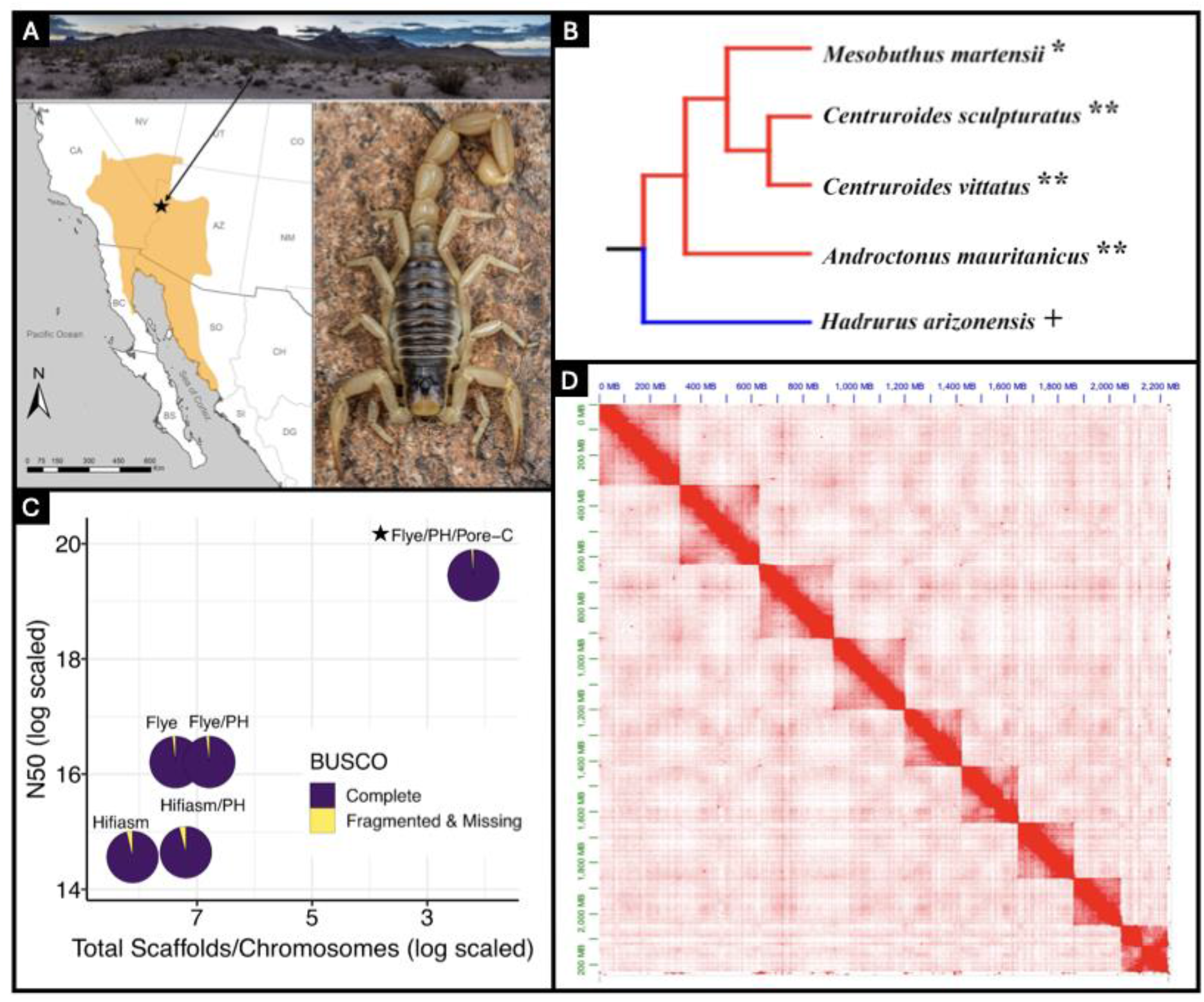
**A)** Range map of *H. arizonensis* with a point (star) denoting the approximate sampling location. The arrow extends to a photo of this region (near Oatman, Arizona, USA). The image on the right is a photo of *H. arizonensis* [both images are courtesy of Brent E. Hendrixson]. **B)** Modified phylogeny from Santibáñez López et al. 2023, including only species with genomes currently available on NCBI. Symbols appended to the species name indicate assembly status: ‘*’ Scaffold level, ‘**’ Contig level, ‘+’ Complete chromosome-scale assembly. The red segments indicate parvorder Buthida while the blue segments indicate parvorder Iurida. **C)** Contiguity and completeness of genome assembly approaches for *H. arizonensis*. These include Flye and Hifiasm, with and without Purge Haplotigs (PH). The final chromosome-scale assembly is denoted with a star. Each assembly is represented by a pie chart to visualize BUSCO completeness. Dark blue indicates complete BUSCOs while yellow indicates missing and/or fragmented BUSCOs. **D)** Chromatin contact map generated from Pore-C data shows the nine chromosomes (2n = 18) that represent 99.56% of the assembled *H. arizonensis* genome.

## Results & Discussion

### Sequencing

Nanopore sequencing generated 13,570,836 high quality long reads, of which 92.8% were sheared (read N50: 14,664 bp) and 7.2% were ultra-long (read N50: 52,738 bp). Following contaminant screening, 63,023 reads (0.46%) were removed (Table S1). The k-mer based genome size estimation was 2.31 Gb from 109.96 Gb of read data (47.6x), double the 1.1 Gb GoaT estimation (Challis *et al*. 2023) (Table S1).

### Genome Assembly

Flye and Hifiasm, with and without Purge Haplotigs (PH), were used to assemble the first genome (Figure 1C; Table S1). The assemblies ranged from 2.2 Gb to 2.6 Gb in length. Flye had an N50 of 10.9 Mb with 1.6 K contigs, and a BUSCO score of C:98.1%, D:3.7%, and F&M:1.9%. Hifiasm had an N50 of 2.1 Mb with 3.4 K contigs, and a BUSCO score of C:96.3%, D:12.2%, and F&M:3.7%. Hifiasm PH had an N50 of 2.3 Mb and 1.3 K contigs, with a BUSCO score of C:95.8%, D:6.2%, and F&M:4.2%. Flye PH had the highest N50 of 10.97 Mb and the fewest contigs (885 in total) (Table S1). Flye PH had a BUSCO score of C:98.1%, D:3.7%, and F&M:1.9%. Flye PH, with the greatest contiguity and completeness, was selected for scaffolding. Contigs less than 3 Kb were removed from Flye PH prior to scaffolding. The final filtered Flye PH assembly had an N50 of 10.9 Mb, 850 contigs, a BUSCO score of C:98.2% [S:94.6%, D:3.6%], F&M:1.8%, and a Merqury QV score of 41.27.

### Genome Scaffolding

The sequenced Pore-C reads yielded 86.9 Gb (∼37x) of data with a mean read quality of 19.8 and read length N50 of 4.3 Kb (Table S2). The wf-pore-c and YaHS pipeline re-oriented all 850 contigs into 242 scaffolds (Figure 1D). Of these, 9 represented 99.56% of the genome, while the remaining 233 accounted for 0.44%. The final chromosome-level assembly was 2.23 Gb in size and had an N50 of 280 Mb, BUSCO score of C:98.6%, D:3.4%, and F&M:1.4%, and Merqury QV of 41.28. While karyotypes of *Hadrurus hirsutus* within Hadruridae suggest an approximate haploid number of 50 (Wilson 1931), scorpion haploid numbers vary significantly, ranging between 5 and 90 (Šťáhlavský *et al*. 2021).

### Repeat Annotation

Prior to structural annotation, RepeatMasker softmasked 1.22 Gb (54.69%) of the genome. Retrotransposons were the most abundant transposable element class in the *H. arizonensis* genome, comprising 29.86% of the repetitive elements identified (Figure 2A). Long interspersed nuclear elements (LINEs) represented the majority of retrotransposons (29.39%), in stark contrast with short interspersed nuclear elements (SINEs) (0.16%) and long terminal repeat elements (LTRs) (0.31%) (Table S3). Interestingly, Bovine-B (RTE/Bov-B) was the most abundant LINE (27.07%). Bov-B elements have a widespread and patchy distribution in eukaryotes and phylogenetic analysis of these elements has identified potential horizontal transfer (HT) vectors in Arthropoda (Ivancevic *et al*. 2018). Approximately 13.57% of the genomes’ repeat elements consist of DNA transposons, with Tc1-IS630-Pogo accounting for 9.77%. This family has been identified in several species, including *Drosophila*, plants, even vertebrates (Gao *et al*. 2020) and in some instances, greatly expanded (Marburger *et al*. 2018).

**Figure 2.**
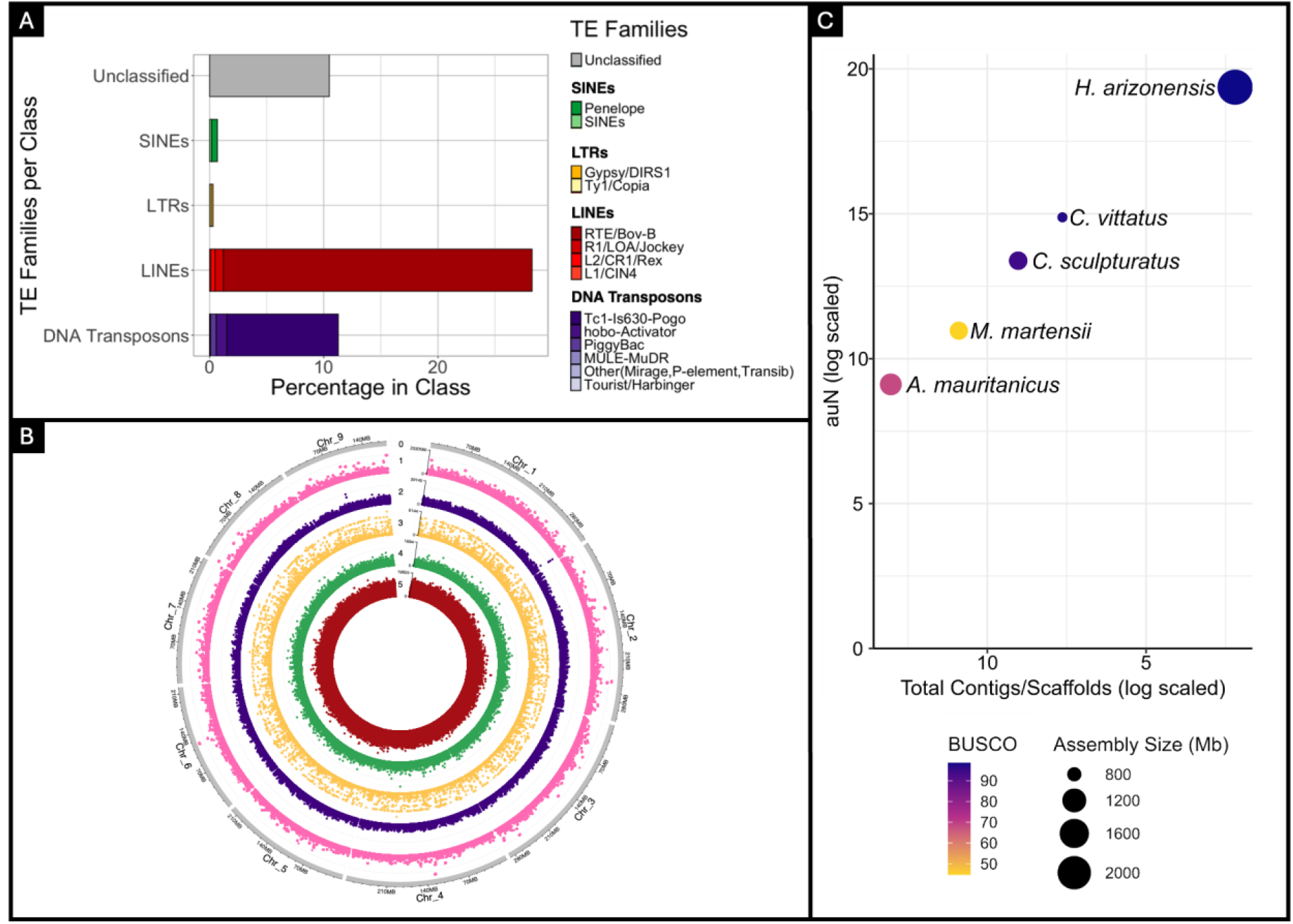
**A)** Percentage of TE families (by repeat class) in the *H. arizonensis* genome assembly. Color gradients were used to differentiate the primary TE families: Unclassified (gray), SINEs (green), LTRs (yellow), LINEs (red), and DNA Transposons (purple). **B)** ShinyCircos (v2.0) representation of the chromosome-scale assembly with the following numbered tracks (10 Kb windows) from outside to inside: (0) primary chromosomes (gray), (1) gene density (pink), (2) DNA transposon density (purple), (3) LTR density (yellow), (4) SINE density (green) and (5) LINE density (red). **C)** Contiguity and completeness of the four scorpion genomes available in NCBI, in addition to *H. arizonensis*. Color gradient represents BUSCO completeness while the size of each point represents the assembly size (Table S7).

### Protein-coding Genes

Six *H. arizonensis* RNA libraries exported from NCBI had an average mapping rate of 92.44% (Table S4). Combined, the filtered EASEL and GeneMark structural annotation yielded 31,841 genes and 90,343 transcripts. These numbers were reduced to 18,996 and 75,256, respectively, following PFAM filtration. With alternative transcripts present, the mono:multi exonic ratio was 0.14 and BUSCO completeness was 97.2% [S:9.7%, D:87.5%] (Figure 2B). After taking the longest isoform, BUSCO completeness decreased to 91.6% [S:87.6%, D:4.0%], while the mono:multi exonic ratio increased to 0.14 (Table S5). The reciprocal BLAST annotation rate of the longest isoform proteins against the RefSeq database was 59.21%. In comparison, the annotation rate for gene family assignment using the EggNOG v4 database was 74.33%. Combined, the final annotation rate for the extracted longest isoforms reached 75.94% (Table S6). While 18,996 genes may be an underestimation, in comparison to the 24,591 predicted protein-coding genes in the *Centruroides sculpturatus* genome (GCF_000671375.1; Schwager *et al*. 2017) they are complete and in range with Araneae (Thomas *et al*. 2020).

### Scorpion Genomes

Presently, only 12 of 237 arachnid genomes hosted at NCBI are chromosome-scale. Prior to assembling *H. arizonensis*, there were no chromosome-scale genome assemblies within the order Scorpiones. As such, this high-quality reference represents the first, with a completeness and contiguity that surpasses the scorpion assemblies available to date (Figure 2C, Figure 1B).

## Materials & Methods

### Collection

One adult female *Hadrurus arizonensis* individual was collected near Oatman, Arizona, USA on August 25th, 2022, and preserved in 100% ethanol (Figure 1A). The legs and pedipalps were removed, rinsed in nuclease-free water, and frozen in liquid nitrogen. The tissue was ground in liquid nitrogen to a fine powder and stored at -80 °C.

### DNA Extraction

*Hadrurus arizonensis* DNA was extracted using the Monarch^Ⓡ^High Molecular Weight (HMW) DNA Extraction Kit: Tissue Protocol (NEB, T3060). Changes made to the protocol include: adding 580 μL of HMW gDNA Tissue Lysis Buffer to approximately 30 mg of frozen ground tissue, lysis incubation was at 56 °C for 45 minutes at 700 rpm and the binding of the gDNA to beads was performed on a vertical rotating mixer at 8 rpm rather than 10 rpm. DNA concentration was measured using a Qubit fluorometer. The gDNA for the standard library was sheared using a Covaris^Ⓡ^G-tube to approximately 25 Kb and small fragments were removed using PacBio’s Short Read Eliminator XS kit. For the ultra-long library, three separate Monarch^Ⓡ^extractions were pooled and eluted in Oxford Nanopore’s EEB buffer.

### Genomic Library Preparation

Oxford Nanopore Technologies^Ⓡ^ Ligation Sequencing Kit V14 (SQK-LSK114) was used to prepare the library on the extracted HMW DNA. DNA repair was performed at 20 °C for 20 minutes followed by 10 minutes at 65 °C to inactivate the enzymes. DNA was eluted at 37 °C. The final library was quantified using a Qubit fluorometer. The flow cell was loaded 3 times each with 10.5 fmol of library. For ultra-long genomic library preparation, the Ultra-Long DNA Kit V14 (SQK-ULK114) was used on the pooled high molecular weight DNA. Both PromethION flow cells ran for 72 hours.

### Pore-C Library Preparation and Sequencing

A total of 150 mg of ground tissue was used for cell crosslinking. Crosslinked cells were resuspended in permeabilization solution and incubated on ice. The chromatin was denatured and the permeabilized cells were digested with restriction enzyme NlaIII. To reverse the crosslinking of the ligated chromatin, the sample suspension was incubated in a thermomixer with periodic rotation. Full details on the library preparation are available (File S1). A total of 1.5 μg of purified DNA was used as input material for the SQK-LSK114 (Oxford Nanopore Technologies, UK) kit and protocol. DNA repair was performed at 20 °C for 20 minutes, followed by 65 °C for 10 minutes to inactivate the enzymes. The PromethION flow cell was run for 96 hours.

### Genome Assembly and Scaffolding

Ultra-long and sheared ONT raw reads were basecalled by Dorado (v7.0.8) and assessed with NanoPlot (v1.33.0) (De Coster & Rademakers 2023). Reads passing a quality threshold (Q > 10) underwent contaminant filtering using Centrifuge (v1.0.4-beta), with a minimum match length of 50 bp against NCBI’s RefSeq bacteria, archaea, and fungi databases (Kim *et al*. 2016). Classified reads were removed, and NanoPlot was rerun. Genome size and coverage was estimated using kmerfreq (v4.0) and GCE (v1.0.2) at k-mer size 21 (Liu *et al*. 2013; Wang *et al*. 2020).

Two long-read *de novo* assembly tools, Flye (v2.8.1) and Hifiasm (v0.19.6-r595), were assessed (Kolmogorov *et al*. 2019; Cheng *et al*. 2021). Flye ran with coverage set to 60, while Hifiasm ran with default parameters. Assembly completeness, contiguity and accuracy were estimated at each assembly stage with BUSCO (v5.4.5) using the arthropoda_odb10 lineage database, Merqury (v1.3), and QUAST (v5.2.0), respectively (Manni *et al*. 2021; Rhie *et al*. 2020; Mikheenko *et al*. 2018). Both assemblies were polished with Medaka (v1.9.1) using model r1041_e82_400bps_sup_g615 and minimap2 (v2.26) aligned raw reads, but neither moved forward due to decreased completeness and quality (*medaka*; Li 2018). To reduce duplication, Purge Haplotigs (v1.1.2) was run on the original assemblies (Roach *et al*. 2018). Low, medium and high read-depth thresholds were set to 9, 26, and 195 in Flye, respectively, and 3, 32, and 195 in Hifiasm. The purged Flye assembly was selected for scaffolding due to superior BUSCO completeness, N50 (contiguity), and accuracy (Merqury QV).

A single Pore-C library was basecalled by Dorado (v7.1.4) from a 96-hour run and used to scaffold the Flye PH assembly. The wf-pore-c (v1.0.0) Nextflow workflow was employed to pre-process and align Pore-C raw reads to the draft genome with Fastcat (v0.14.1) and minimap2 (v2.26-r1175) respectively (*wf-pore-c*). The resulting alignments were converted into a 4DN-format pairs file with pairtools parse2 (v1.0.2) (Open2C *et al*. 2023). YaHS (v1.1) conducted multiple rounds of scaffolding using the pairs file format, genome and ‘GATG’ restriction enzyme flag. The resulting Pore-C alignment and APG file were run through juicer (v1.2) pre and juicer_tools (v1.9.9) for visualization with Juicebox (Zhou *et al*. 2023; Durand *et al*. 2016a; Durand *et al*. 2016b). The 9 manually curated chromosomes were assessed with BUSCO, Merqury, and QUAST.

### Genome Annotation

Transposable elements were identified with RepeatModeler (v2.02) and the reference was softmasked with RepeatMasker (v4.1.4) (Flynn *et al*. 2020; Smit *et al*. 2015). Six *H. arizonensis* RNA libraries were imported from NCBI (PRJNA340270) and used in the EASEL (v1.5) pipeline along with the softmasked draft genome and arthropoda OrthoDB (v11) protein sequences to predict protein-coding genes (Webster *et al*. n.d.). The invertebrate training set was configured to filter false-positive predictions. The same inputs were utilized in the GeneMark-ETP (Bruna *et al*. 2024) gene prediction tool, which is part of the BRAKER3 (v3.0.2) pipeline (Gabriel *et al*. 2024). EASEL (filtered) and GeneMark (hmm) genes were independently mapped to the Pore-C chromosome-level assembly with Liftoff (v1.6.3) (Shumate *et al*. 2021). The resulting GFF files were combined using the AGAT (v1.2) toolkit (Dainat *et al*. 2022). Protein sequences were extracted and scanned for protein domains using Pfam-A.hmm (v3.1b2) and HMMER (v3.3.2) (Mistry *et al*. 2021; Eddy *et al*. 2011). Sequences without a domain were removed. Final summary statistics, including BUSCO, were run on the filtered proteins. Furthermore, EnTAP (v1.0.1) was run with the complete RefSeq database (v208) at 70/70 coverage to functionally annotate the predicted proteins (Hart *et al*. 2020). Summary statistics, BUSCO, and EnTAP were also run on the longest isoforms to represent the unique gene space.

## Data Availability Statement

The nanopore long-reads used for the genome assembly have been deposited under the NCBI Bioproject PRJNA1072625. This references SAMN39896521 (standard nanopore) and SAMN39747025 (ultra-long nanopore). The assembly and annotation are pending approval at NCBI, and are hosted, with all project code on the following Gitlab: https://gitlab.com/PlantGenomicsLab/hadrurus-arizonensis-genome-assembly-and-annotation.

## Acknowledgments

The authors acknowledge contributions of the HPC resources from the Computational Biology Core within the Institute for Systems Genomics.

