## Supplemental File 1 for "Unveiling the genetic blueprint of a desert scorpion: A chromosome-level genome of *Hadrurus arizonensis* provides the first reference for Parvorder Iurida"

### *Pore-C library preparation methods*

*Cell crosslinking:*

150mg of ground tissue was washed in cold PBS, crosslinked in 1% formaldehyde, and incubated for 10 minutes at room temperature. The reaction was quenched with 125mM glycine. The cell suspension was incubated for five minutes at room temperature, followed by 10 minutes on ice. Fixed cells were pelleted at 500 × g for 5 minutes at 4 °C and washed twice with 5 ml cold 1× PBS. The cell pellet was stored at −80 °C until further processing.

*Chromatin digestion and ligation:*

Crosslinked cells were resuspended in 550 µL permeabilization solution and incubated on ice for 15 minutes with regular inversion. Nuclei were pelleted at 4°C for 10 min at 500 × g, and the supernatant was discarded. The pellet was washed in 200 µL of 1.5x CutSmart buffer (NEB) and re-pelleted at 500 x g for 5 minutes at 4°C. The pellet was resuspended in 300ul of 1.5x CutSmart buffer (NEB). The chromatin was denatured by adding 0.1% SDS (final concentration) and incubating in a thermomixer at 300 rpm at 65°C for 10 minutes. The sample was chilled on ice prior to adding 10% (v/v) Triton X-100 for a final concentration of 1% and incubated on ice for 10 min. The permeabilized cells were digested with restriction enzyme NlaIII for 18 hours at 37°C. The restriction enzyme was denatured via incubation in a thermomixer at 65°C with 300 rpm rotation for 20 minutes. The sample was cooled to room temperature.

The proximity ligation was incubated for 6 hours at 16°C with intermittent mixing. The reaction was stopped by adding 0.1% SDS (final concentration) and incubating in a thermomixer at 300 rpm at 65°C for 20 minutes.

*Protein degradation and DNA purification:*

To reverse the crosslinking of the ligated chromatin, the sample suspension was incubated in a thermomixer at 56°C for 18 hours with periodic rotation. The sample was incubated on ice with an equal volume of 25:24:1 phenol:chloroform:isoamyl alcohol and centrifuged at 16,000 g at 15°C for 15 minutes. DNA precipitation was performed by adding 0.1 volumes of 3 M sodium acetate and 3 volumes 100% ethanol to the upper aqueous solution. DNA was resuspended in 75 µl of TE buffer and stored at 4°C.
