## Supplemental Tables 1-7 for "Unveiling the genetic blueprint of a desert scorpion: A chromosome-level genome of *Hadrurus arizonensis* provides the first reference for Parvorder Iurida"

**Table S1.** Genome Size, Raw Read Statistics, & Initial Assembly Statistics

**Genome Size Estimate:**

| | | | |
|---|---|---|---|
| kmerfreq+GCE: | Centrifuge Filtered | | |
| Sheared, Filtered | 2.32818 Gbp | | |
| Ultra Long, Filtered | 2.45343 Gbp | | |
| Shear+UL, Filtered | 2.31133 Gbp | | |
| GoaT | 1.1G | | |
| **Raw Reads:** | Shear | | |
| | ALL (PASS + FAIL) | PASS | FAIL |
| Active channels: | 2,706.00 | 2,664.00 | 2,696.00 |
| Mean read length: | 7,304.80 | 7,297.80 | 7,341.10 |
| Mean read quality: | 17.4 | 19.4 | 6.7 |
| Median read length: | 4,172.00 | 4,141.00 | 4,316.00 |
| Median read quality: | 18.6 | 19.6 | 6.5 |
| Number of reads: | 15,110,803.00 | 12,654,298.00 | 2,456,505.00 |
| Read length N50: | 14,597.00 | 14,664.00 | 14,252.00 |
| STDEV read length: | 8,352.00 | 8,161.60 | 9,270.90 |
| Total bases: | 110,381,410,046.00 | 92,348,047,592.00 | 18,033,362,454.00 |
| Number, percentage and megabases of reads above quality cutoffs | | | |
| >Q5 | 14847728 (98.3%) 108399.0Mb | 12654298 (100.0%) 92348.0Mb | 2193430 (89.3%) 16051.0Mb |
| >Q7 | 13435214 (88.9%) 97469.1Mb | 12654298 (100.0%) 92348.0Mb | 780916 (31.8%) 5121.0Mb |
| >Q10 | 12654297 (83.7%) 92348.0Mb | 12654297 (100.0%) 92348.0Mb | 0 (0.0%) 0.0Mb |
| >Q12 | 12218609 (80.9%) 89245.0Mb | 12218609 (96.6%) 89245.0Mb | 0 (0.0%) 0.0Mb |
| >Q15 | 11044921 (73.1%) 81680.8Mb | 11044921 (87.3%) 81680.8Mb | 0 (0.0%) 0.0Mb |
| Top 5 highest mean basecall quality scores and their read lengths | | | |
| 1 | 44.6 (257) | 44.6 (257) | 10.0 (7849) |
| 2 | 44.1 (173) | 44.1 (173) | 10.0 (22889) |
| 3 | 44.0 (171) | 44.0 (171) | 10.0 (1189) |
| 4 | 44.0 (290) | 44.0 (290) | 10.0 (565) |
| 5 | 44.0 (248) | 44.0 (248) | 10.0 (9706) |
| Top 5 longest reads and their mean basecall quality score | | | |
| 1 | 1089080 (3.7) | 497912 (12.2) | 1089080 (3.7) |
| 2 | 1065876 (3.1) | 496343 (10.7) | 1065876 (3.1) |

| | | | |
|---|---|---|---|
| 3 | 746502 (3.8) | 488058 (10.8) | 746502 (3.8) |
| 4 | 624323 (3.3) | 481137 (11.3) | 624323 (3.3) |
| 5 | 608640 (5.4) | 474741 (10.9) | 608640 (5.4) |

| **Assembly Statistics:** | Flye | Flye + PH | Flye + PH + remove < 3000kb |
|---|---|---|---|
| BUSCO Embryophyta | | | |
| Complete BUSCOs (C) | 993 (98.1%) | 993 (98.1%) | 994 (98.2) |
| Complete and single-copy BUSCOs (S) | 956 (94.4%) | 955 (94.3%) | 958 (94.6) |
| Complete and duplicated BUSCOs (D) | 37 (3.7%) | 38 (3.8%) | 36 (3.6%) |
| Fragmented BUSCOs (F) | 13 (1.3%) | 13 (1.3%) | 12 (1.2%) |
| Missing BUSCOs (M) | 7 (0.6%) | 7 (0.6%) | 7(0.6%) |
| Total BUSCO groups searched | 1013 | 1013 | 1013 |
| Merqury | | | |
| QV | 39.4886 | 37.4648 | 41.2745 |
| Error | 1.13E-04 | 1.79E-04 | 7.46E-05 |
| Quast | | | |
| # contigs | 1,591 | 889 | 850 |
| Largest contig | 50,833,969 | 50,833,625 | 50,833,969 |
| Total length | 2,260,818,358 | 2,238,006,049 | 2,237,111,498 |
| GC % | 32.23 | 32.21 | 32.22 |
| N50 | 10,940,222 | 10,965,598 | 10,965,104 |

| | | | |
|---|---|---|---|
| Ultra Long | | | Centrifuge Filtered |
| ALL (PASS + FAIL) | PASS | FAIL | Shear + Ultra Long |
| 2,413.00 | 2,350.00 | 2,375.00 | N/A |
| 18,585.10 | 19,151.90 | 16,049.50 | 8,102.50 |
| 16.9 | 19.2 | 6.2 | 18.2 |
| 6,215.00 | 6,483.00 | 5,182.00 | 4,283.00 |
| 18.7 | 19.7 | 6.2 | 18.5 |
| 1,198,543.00 | 979,561.00 | 218,982.00 | 13,570,836.00 |
| 51,959.00 | 52,738.00 | 47,833.00 | 16,053.00 |
| 28,573.70 | 28,749.50 | 27,632.10 | 11,279.70 |
| 22,275,048,152.00 | 18,760,498,613.00 | 3,514,549,539.00 | 109,958,199,491.00 |
| 1137348 (94.9%) 21385.3Mb | 979561 (100.0%) 18760.5Mb | 157787 (72.1%) 2624.8Mb | 152 (100.0%) 109958.0Mb |
| 1040593 (86.8%) 19760.3Mb | 979561 (100.0%) 18760.5Mb | 61032 (27.9%) 999.8Mb | 062 (100.0%) 109957.9Mb |
| 979561 (81.7%) 18760.5Mb | 979561 (100.0%) 18760.5Mb | 0 (0.0%) 0.0Mb | 2651 (99.3%) 109830.6Mb |
| 942754 (78.7%) 18145.4Mb | 942754 (96.2%) 18145.4Mb | 0 (0.0%) 0.0Mb | 6678 (95.0%) 106013.1Mb |
| 856364 (71.5%) 16598.6Mb | 856364 (87.4%) 16598.6Mb | 0 (0.0%) 0.0Mb | 15459 (81.2%) 96264.1Mb |
| 38.1 (915) | 38.1 (915) | 10.0 (1064) | 41.0 (259) |
| 38.0 (1283) | 38.0 (1283) | 10.0 (24570) | 41.0 (297) |
| 37.2 (907) | 37.2 (907) | 10.0 (6823) | 40.8 (228) |
| 36.1 (1263) | 36.1 (1263) | 10.0 (36766) | 40.8 (303) |
| 36.0 (965) | 36.0 (965) | 10.0 (10157) | 40.5 (313) |
| 1452070 (3.3) | 376012 (16.3) | 1452070 (3.3) | 497912 (12.2) |
| 1189644 (4.6) | 354472 (18.9) | 1189644 (4.6) | 496343 (10.7) |

| | | | |
|---|---|---|---|
| 1036457 (3.5) | 342052 (15.5) | 1036457 (3.5) | 488058 (10.8) |
| 941963 (3.5) | 341544 (17.7) | 941963 (3.5) | 481137 (11.3) |
| 913558 (2.6) | 329205 (18.6) | 913558 (2.6) | 474741 (10.9) |

| Hifiasm | Hifiasm + PH |
|---|---|
| 976 (96.3%) | 971 (95.8%) |
| 852 (84.1%) | 907 (89.5%) |
| 124 (12.2%) | 64 (6.3%) |
| 20 (2.0%) | 21 (2.1%) |
| 17 (1.7%) | 21 (2.1%) |
| 1013 | 1013 |
| 50.5289 | 39.7118 |
| 8.85E-06 | 1.07E-04 |
| 3,359 | 1627 |
| 11,774,485 | 11786046 |
| 2,569,518,486 | 2,324,022,605 |
| 32.22 | 32.22 |
| 2,109,929 | 2,295,267 |

**Table S2.** Pore-C Scaffolding statistics with NanoPlot, Quast, BUSCO, and Merqury.

**Scaffolding**

| **POREC Raw Reads NanoPlot** | **Passing** |
|---|---|
| Active channels: | 2,640.00 |
| Mean read length: | 1,747.60 |
| Mean read quality: | 19.8 |
| Median read length: | 754 |
| Median read quality: | 19.7 |
| Number of reads: | 49,725,252.00 |
| Read length N50: | 4,226.00 |
| STDEV read length: | 3,107.40 |
| Total bases: | 86,902,330,304.00 |
| Number, percentage and megabases of reads above quality cutoffs | |
| >Q5 | 49725252 (100.0%) 86902.3Mb |
| >Q7 | 49725252 (100.0%) 86902.3Mb |
| >Q10 | 49725251 (100.0%) 86902.3Mb |
| >Q12 | 48283598 (97.1%) 84492.9Mb |
| >Q15 | 43721106 (87.9%) 77992.1Mb |
| Top 5 highest mean basecall quality scores and their read lengths | |
| 1 | 46.9 (110) |
| 2 | 45.5 (115) |
| 3 | 44.9 (68) |
| 4 | 44.8 (141) |
| 5 | 44.5 (250) |
| Top 5 longest reads and their mean basecall quality score | |
| 1 | 250271 (12.7) |
| 2 | 221149 (11.7) |
| 3 | 197851 (11.2) |
| 4 | 189985 (11.2) |
| 5 | 185930 (10.2) |
| Coverage | 37.33 |

| **Scaffold** | **Length (bp)** |
|---|---|
| **scaffold_1** | **315073531** |
| **scaffold_2** | **313324015** |
| **scaffold_3** | **289489910** |
| **scaffold_4** | **279785186** |
| **scaffold_5** | **222232424** |
| **scaffold_6** | **221366528** |
| **scaffold_7** | **219092046** |
| **scaffold_8** | **186296334** |
| **scaffold_9** | **180654565** |
| scaffold_10 | 1273507 |
| scaffold_11 | 1180150 |
| scaffold_12 | 1127565 |
| scaffold_13 | 553832 |
| scaffold_14 | 399942 |
| scaffold_15 | 377387 |
| scaffold_16 | 225993 |
| scaffold_17 | 173354 |
| scaffold_18 | 144515 |
| scaffold_19 | 138891 |
| scaffold_20 | 128509 |
| scaffold_21 | 125000 |
| scaffold_22 | 120188 |
| scaffold_23 | 119737 |
| scaffold_24 | 106060 |
| scaffold_25 | 105196 |
| scaffold_26 | 99360 |
| scaffold_27 | 97523 |
| scaffold_28 | 96931 |
| scaffold_29 | 95823 |
| scaffold_30 | 91488 |
| scaffold_31 | 85941 |
| scaffold_32 | 76473 |
| scaffold_33 | 66846 |

| | |
|---|---|
| scaffold_34 | 63000 |
| scaffold_35 | 63000 |
| scaffold_36 | 62727 |
| scaffold_37 | 59883 |
| scaffold_38 | 59596 |
| scaffold_39 | 59198 |
| scaffold_40 | 58699 |
| scaffold_41 | 57821 |
| scaffold_42 | 55764 |
| scaffold_43 | 54000 |
| scaffold_44 | 53560 |
| scaffold_45 | 49827 |
| scaffold_46 | 47000 |
| scaffold_47 | 46186 |
| scaffold_48 | 45180 |
| scaffold_49 | 43487 |
| scaffold_50 | 42137 |
| scaffold_51 | 41088 |
| scaffold_52 | 38243 |
| scaffold_53 | 37835 |
| scaffold_54 | 35702 |
| scaffold_55 | 35197 |
| scaffold_56 | 33183 |
| scaffold_57 | 32291 |
| scaffold_58 | 32202 |
| scaffold_59 | 31429 |
| scaffold_60 | 30999 |
| scaffold_61 | 28166 |
| scaffold_62 | 27992 |
| scaffold_63 | 27678 |
| scaffold_64 | 23057 |
| scaffold_65 | 22126 |
| scaffold_66 | 21499 |
| scaffold_67 | 21082 |
| scaffold_68 | 19846 |

| | |
|---|---|
| scaffold_69 | 19807 |
| scaffold_70 | 19711 |
| scaffold_71 | 19675 |
| scaffold_72 | 19463 |
| scaffold_73 | 19372 |
| scaffold_74 | 18648 |
| scaffold_75 | 18137 |
| scaffold_76 | 17863 |
| scaffold_77 | 17681 |
| scaffold_78 | 17622 |
| scaffold_79 | 17455 |
| scaffold_80 | 17202 |
| scaffold_81 | 17013 |
| scaffold_82 | 16365 |
| scaffold_83 | 16365 |
| scaffold_84 | 16355 |
| scaffold_85 | 16278 |
| scaffold_86 | 16243 |
| scaffold_87 | 16168 |
| scaffold_88 | 16066 |
| scaffold_89 | 16023 |
| scaffold_90 | 15815 |
| scaffold_91 | 15667 |
| scaffold_92 | 15267 |
| scaffold_93 | 15125 |
| scaffold_94 | 14981 |
| scaffold_95 | 14373 |
| scaffold_96 | 14086 |
| scaffold_97 | 14000 |
| scaffold_98 | 14000 |
| scaffold_99 | 13738 |
| scaffold_100 | 13591 |
| scaffold_101 | 13485 |
| scaffold_102 | 13047 |
| scaffold_103 | 12910 |

| | |
|---|---|
| scaffold_104 | 12679 |
| scaffold_105 | 12643 |
| scaffold_106 | 12531 |
| scaffold_107 | 12251 |
| scaffold_108 | 12080 |
| scaffold_109 | 12000 |
| scaffold_110 | 12000 |
| scaffold_111 | 12000 |
| scaffold_112 | 11819 |
| scaffold_113 | 11781 |
| scaffold_114 | 11749 |
| scaffold_115 | 11408 |
| scaffold_116 | 11389 |
| scaffold_117 | 11278 |
| scaffold_118 | 11000 |
| scaffold_119 | 10905 |
| scaffold_120 | 10903 |
| scaffold_121 | 10893 |
| scaffold_122 | 10763 |
| scaffold_123 | 10720 |
| scaffold_124 | 10398 |
| scaffold_125 | 9997 |
| scaffold_126 | 9921 |
| scaffold_127 | 9850 |
| scaffold_128 | 9385 |
| scaffold_129 | 9213 |
| scaffold_130 | 9181 |
| scaffold_131 | 9000 |
| scaffold_132 | 9000 |
| scaffold_133 | 8972 |
| scaffold_134 | 8968 |
| scaffold_135 | 8939 |
| scaffold_136 | 8938 |
| scaffold_137 | 8784 |
| scaffold_138 | 8691 |

| | |
|---|---|
| scaffold_139 | 8372 |
| scaffold_140 | 8335 |
| scaffold_141 | 8247 |
| scaffold_142 | 8000 |
| scaffold_143 | 8000 |
| scaffold_144 | 7923 |
| scaffold_145 | 7871 |
| scaffold_146 | 7852 |
| scaffold_147 | 7834 |
| scaffold_148 | 7657 |
| scaffold_149 | 7534 |
| scaffold_150 | 7437 |
| scaffold_151 | 7173 |
| scaffold_152 | 7000 |
| scaffold_153 | 7000 |
| scaffold_154 | 6744 |
| scaffold_155 | 6725 |
| scaffold_156 | 6716 |
| scaffold_157 | 6627 |
| scaffold_158 | 6569 |
| scaffold_159 | 6529 |
| scaffold_160 | 6510 |
| scaffold_161 | 6386 |
| scaffold_162 | 6328 |
| scaffold_163 | 6300 |
| scaffold_164 | 6266 |
| scaffold_165 | 6160 |
| scaffold_166 | 6151 |
| scaffold_167 | 6103 |
| scaffold_168 | 6000 |
| scaffold_169 | 6000 |
| scaffold_170 | 5688 |
| scaffold_171 | 5637 |
| scaffold_172 | 5257 |
| scaffold_173 | 5159 |

| | |
|---|---|
| scaffold_174 | 5147 |
| scaffold_175 | 5147 |
| scaffold_176 | 5142 |
| scaffold_177 | 5027 |
| scaffold_178 | 5019 |
| scaffold_179 | 5000 |
| scaffold_180 | 5000 |
| scaffold_181 | 4989 |
| scaffold_182 | 4928 |
| scaffold_183 | 4919 |
| scaffold_184 | 4842 |
| scaffold_185 | 4775 |
| scaffold_186 | 4764 |
| scaffold_187 | 4763 |
| scaffold_188 | 4702 |
| scaffold_189 | 4674 |
| scaffold_190 | 4634 |
| scaffold_191 | 4526 |
| scaffold_192 | 4367 |
| scaffold_193 | 4278 |
| scaffold_194 | 4107 |
| scaffold_195 | 4044 |
| scaffold_196 | 4000 |
| scaffold_197 | 3927 |
| scaffold_198 | 3710 |
| scaffold_199 | 3539 |
| scaffold_200 | 3509 |
| scaffold_201 | 3433 |
| scaffold_202 | 3395 |
| scaffold_203 | 3356 |
| scaffold_204 | 3340 |
| scaffold_205 | 3277 |
| scaffold_206 | 3145 |
| scaffold_207 | 3143 |
| scaffold_208 | 3084 |

| | |
|---|---|
| scaffold_209 | 3000 |
| scaffold_210 | 3000 |
| scaffold_211 | 3000 |
| scaffold_212 | 3000 |
| scaffold_213 | 3000 |
| scaffold_214 | 3000 |
| scaffold_215 | 3000 |
| scaffold_216 | 3000 |
| scaffold_217 | 3000 |
| scaffold_218 | 3000 |
| scaffold_219 | 2000 |
| scaffold_220 | 2000 |
| scaffold_221 | 2000 |
| scaffold_222 | 2000 |
| scaffold_223 | 2000 |
| scaffold_224 | 2000 |
| scaffold_225 | 2000 |
| scaffold_226 | 2000 |
| scaffold_227 | 2000 |
| scaffold_228 | 2000 |
| scaffold_229 | 1000 |
| scaffold_230 | 1000 |
| scaffold_231 | 1000 |
| scaffold_232 | 1000 |
| scaffold_233 | 1000 |
| scaffold_234 | 1000 |
| scaffold_235 | 1000 |
| scaffold_236 | 1000 |
| scaffold_237 | 1000 |
| scaffold_238 | 1000 |
| scaffold_239 | 1000 |
| scaffold_240 | 1000 |
| scaffold_241 | 1000 |
| scaffold_242 | 1000 |

| | |
|---|---|
| **QUAST** | |
| # contigs (>= 0 bp) | 9 |
| # contigs (>= 1000 bp) | 9 |
| # contigs (>= 5000 bp) | 9 |
| # contigs (>= 10000 bp) | 9 |
| # contigs (>= 25000 bp) | 9 |
| # contigs (>= 50000 bp) | 9 |
| Total length (>= 0 bp) | 22273145 |
| Total length (>= 1000 bp) | 22273145 |
| Total length (>= 5000 bp) | 22273145 |
| Total length (>= 10000 bp) | 22273145 |
| Total length (>= 25000 bp) | 22273145 |
| **Total length (>= 50000 bp)** | **22273145** |
| # contigs | 9 |
| **Largest contig** | **31507353** |
| Total length | 22273145 |
| GC (%) | 32.22 |
| **N50** | **27978518** |
| N90 | 18629633 |
| auN | 25737789 |
| L50 | 4 |
| **L90** | **8** |
| # N's per 100 kbp | 3.48 |
| **BUSCO** | |
| | C:98.6%[S:95.2%,D:3.4%],F:1.0%,M:0.4%,n:1013 |
| Complete BUSCOs (C) | 998 |
| Complete and single-copy BUSCOs (S) | 964 |
| **Complete and duplicated BUSCOs (D)** | 34 |
| Fragmented BUSCOs (F) | 10 |
| Missing BUSCOs (M) | 5 |
| Total BUSCO groups searched | 1013 |
| **MERQURY** | |
| **QV** | 41.2751 |

| | |
|---|---|
| Error | 7.46E-05 |

0.121991[illegible]

**Table S3.** Repeat elements statistics after RepeatMasker

**Repeats**

file name: H_arizonensis_chr_unmasked.fasta
sequences: 9
total length: 2227314539 bp (2227237039 bp excl N/X-runs)
GC level: 32.22 %
bases masked: 1218188793 bp ( 54.69 %)

| | Number of elements | Length occpupied of sequences | Percentage of sequence |
|---|---|---|---|
| **Retroelements:** | 906049 | 665117944 bp | 29.86 % |
| SINEs | 22371 | 3475099 bp | 0.16% |
| Penelope | 45183 | 11649211 bp | 0.52% |
| LINEs | 872874 | 654707858 bp | 29.39 % |
| CRE/SLACS | 0 | 0 bp | 0% |
| L2/CR1/Rex | 18266 | 8283369 bp | 0.37% |
| R1/LOA/Jockey | 17285 | 16174748 bp | 0.73 % |
| R2/R4/NeSL | 0 | 0 bp | 0% |
| RTE/Bov-B | 763412 | 602887460 bp | 27.07 % |
| L1/CIN4 | 4322 | 2330710 bp | 0.10 % |
| **LTR elements:** | 10804 | 6934987 bp | 0.31 % |
| BEL/Pao | 0 | 0 bp | 0% |
| Ty1/Copia | 2585 | 3219682 bp | 0.14 % |
| Gypsy/DIRS1 | 6044 | 3466071 bp | 0.16 % |
| Retroviral | 0 | 0 bp | 0% |
| **DNA transposons:** | 677464 | 302252149 bp | 13.57 % |
| hobo-Activator | 52099 | 20868774 bp | 0.94 % |
| Tc1-IS630-Pogo | 527054 | 217709931 bp | 9.77 % |
| En-Spm | 0 | 0 bp | 0% |
| MULE-MuDR | 3117 | 405809 bp | 0.02 % |
| PiggyBac | 25277 | 10813876 bp | 0.49 % |
| Tourist/Harbinger | 651 | 277372 bp | 0.01% |
| Other (Mirage, P-element, Transib) | 1577 | 1126778 bp | 0.05 % |

| | | | |
|---|---|---|---|
| Rolling-circles | 11641 | 4128768 bp | 0.19 % |
| Unclassified: | 1083252 | 233584851 bp | 10.49 % |
| Total interspersed repeats: | | 1200954944 bp | 53.92 % |
| Small RNA: | 5478 | 986345 bp | 0.04 % |
| Satellites: | 0 | 0 bp | 0.00 % |
| Simple repeats: | 248241 | 10558201 bp | 0.47 % |
| Low complexity: | 46012 | 2210322 bp | 0.10 % |

**Table S4.** NCBI RNA Libraries alignment rates to the genome and quality control results

| **NCBI Accessions:** | **SRR24872962** | **SRR24872963** | **SRR24872964** |
|---|---|---|---|
| **Alignment Rates:** | 93.17 | 93.34 | 92.78 |
| **Quality Control:** | fastp version: | 0.23.2 | |
| **General** | paired end (150 cycles + 150 cycles) | paired end (150 cycles + 150 cycles) | paired end (150 cycles + 150 cycles) |
| mean length before filtering: | 150bp, 150bp | 150bp, 150bp | 150bp, 150bp |
| mean length after filtering: | 138bp, 138bp | 141bp, 141bp | 141bp, 141bp |
| duplication rate: | 24.49% | 32.19% | 15.95% |
| Insert size peak: | 150 | 150 | 146 |
| **Before filtering** | | | |
| total reads: | 26.042326 M | 27.626284 M | 44.809686 M |
| total bases: | 3.906349 G | 4.143943 G | 6.721453 G |
| Q20 bases: | 3.817206 G (97.718004%) | 4.052564 G (97.794880%) | 6.205631 G (92.325734%) |
| Q30 bases: | 3.674891 G (94.074829%) | 3.899643 G (94.104648%) | 5.885195 G (87.558383%) |
| GC content: | 36.68% | 34.80% | 37.50% |
| **After filtering** | | | |
| total reads: | 25.785774 M | 27.314118 M | 44.341840 M |
| total bases: | 3.575646 G | 3.863448 G | 6.285138 G |
| Q20 bases: | 3.513211 G (98.253889%) | 3.796168 G (98.258557%) | 5.863249 G (93.287519%) |
| Q30 bases: | 3.390204 G (94.813756%) | 3.660358 G (94.743315%) | 5.573877 G (88.683455%) |
| GC content: | 35.56% | 33.85% | 36.69% |
| **Filtering result** | | | |
| reads passed filters: | 25.785774 M (99.014865%) | 27.314118 M (98.870040%) | 44.341840 M (98.955927%) |
| reads with low quality: | 251.230000 K (0.964699%) | 306.852000 K (1.110725%) | 446.840000 K (0.997195%) |
| reads with too many N: | 5.322000 K (0.020436%) | 5.314000 K (0.019235%) | 21.006000 K (0.046878%) |
| reads too short: | 0 (0.000000%) | 0 (0.000000%) | 0 (0.000000%) |

| SRR24872965 | SRR24872966 | SRR24872967 |
|---|---|---|
| 92.79 | 89.87 | 92.69 |
| paired end (150 cycles + 150 cycles) | paired end (150 cycles + 150 cycles) | paired end (150 cycles + 150 cycles) |
| 150bp, 150bp | 150bp, 150bp | 150bp, 150bp |
| 145bp, 145bp | 143bp, 143bp | 143bp, 143bp |
| 9.29% | 10.30% | 8.77% |
| 150 | 150 | 155 |
| 51.450696 M | 33.886252 M | 44.771784 M |
| 7.717604 G | 5.082938 G | 6.715767600 G |
| 7.166948 G (92.864932%) | 4.639877 G (91.283364%) | 6.238024197 G (92.8862%) |
| 6.802414 G (88.141529%) | 4.381040 G (86.191093%) | 5.924806044 G (88.2223%) |
| 36.97% | 37.45% | 36.75% |
| 51.184946 M | 33.164022 M | 44.552338 M |
| 7.463272 G | 4.768901 G | 6.381915 G |
| 6.960525 G (93.263708%) | 4.416806 G (92.616846%) | 5.962306 G (93.425025%) |
| 6.613520 G (88.614211%) | 4.181664 G (87.686120%) | 5.671792 G (88.872888%) |
| 36.61% | 36.78% | 36.08% |
| 51.184946 M (99.483486%) | 33.164022 M (97.868664%) | 44.552338 M (99.509856%) |
| 240.398000 K (0.467240%) | 706.450000 K (2.084769%) | 196.888000 K (0.439759%) |
| 25.352000 K (0.049274%) | 15.780000 K (0.046568%) | 22.558000 K (0.050384%) |
| 0 (0.000000%) | 0 (0.000000%) | 0 (0.000000%) |

**Table S5.** Structural annotation statistics before and after extracting the longest isoforms of each gene with AGA[illegible]

| Structural Annotation | Alternative Transcripts | Longest Isoform |
|---|---|---|
| **Summary Statistics (AGAT)** | | |
| Number of gene | 18,996 | 18996 |
| Number of transcript | 75,256 | 18996 |
| Number of mrnas with utr both sides | 14954 | 3036 |
| Number of mrnas with at least one utr | 15,006 | 3047 |
| Number of cds | 75,256 | 18996 |
| Number of exon | 908005 | 156265 |
| Number of five_prime_utr | 14966 | 3037 |
| Number of start_codon | 61499 | 9342 |
| Number of stop_codon | 61490 | 9340 |
| Number of three_prime_utr | 14994 | 3046 |
| Number of exon in cds | 881250 | 154644 |
| Number of exon in five_prime_utr | 33553 | 4396 |
| Number of exon in three_prime_utr | 22949 | 3283 |
| Number of intron in cds | 805994 | 135648 |
| Number of intron in exon | 832749 | 137269 |
| Number of intron in five_prime_utr | 18,587 | 1359 |
| Number of intron in three_prime_utr | 7,955 | 237 |
| Number gene overlapping | 1836 | 1130 |
| Number of single exon gene | 2,255 | 2320 |
| Mono:multi-exonic ratio | 0.135 | 0.139 |
| Number of single exon transcript | 3,669 | 2320 |
| mean transcripts per gene | 4 | 1 |
| mean cdss per transcript | 1 | 1 |
| mean exons per transcript | 12.1 | 8.2 |
| mean five_prime_utrs per transcript | 0.2 | 0.2 |
| mean start_codons per transcript | 0.8 | 0.5 |
| mean stop_codons per transcript | 0.8 | 0.5 |
| mean three_prime_utrs per transcript | 0.2 | 0.2 |
| mean exons per cds | 11.7 | 8.1 |
| mean exons per five_prime_utr | 2.2 | 1.4 |
| mean exons per three_prime_utr | 1.5 | 1.1 |
| mean introns in cdss per transcript | 10.7 | 7.1 |

| | | |
|---|---|---|
| mean introns in exons per transcript | 11.1 | 7.2 |
| mean introns in five_prime_utrs per transcri | 0.2 | 0.1 |
| mean introns in three_prime_utrs per transc | 0.1 | 0 |
| Total gene length (bp) | 1720277191 | 1720277191 |
| Total transcript length (bp) | 9691453464 | 1419893230 |
| Total cds length (bp) | 150900803 | 30385340 |
| Total exon length (bp) | 177241546 | 35329971 |
| Total five_prime_utr length (bp) | 5837879 | 725533 |
| Total start_codon length (bp) | 184497 | 28026 |
| Total stop_codon length (bp) | 184470 | 28020 |
| Total three_prime_utr length (bp) | 20502864 | 4219098 |
| Total intron length per cds (bp) | 9151902898 | 1356812033 |
| Total intron length per exon (bp) | 9514211918 | 1384563259 |
| Total intron length per five_prime_utr (bp) | 284521341 | 24049931 |
| Total intron length per three_prime_utr (bp) | 74841236 | 3246639 |
| mean gene length (bp) | 90559 | 90559 |
| mean transcript length (bp) | 128779 | 74746 |
| mean cds length (bp) | 2005 | 1599 |
| mean exon length (bp) | 195 | 226 |
| mean five_prime_utr length (bp) | 390 | 238 |
| mean start_codon length (bp) | 3 | 3 |
| mean stop_codon length (bp) | 3 | 3 |
| mean three_prime_utr length (bp) | 1367 | 1385 |
| mean cds piece length (bp) | 171 | 196 |
| mean five_prime_utr piece length (bp) | 173 | 165 |
| mean three_prime_utr piece length (bp) | 893 | 1285 |
| mean intron in cds length (bp) | 11354 | 10002 |
| mean intron in exon length (bp) | 11425 | 10086 |
| mean intron in five_prime_utr length (bp) | 15307 | 17696 |
| mean intron in three_prime_utr length (bp) | 9408 | 13698 |
| Longest gene (bp) | 2089961 | 2089961 |
| Longest transcript (bp) | 1636570 | 1636570 |
| Longest cds (bp) | 64446 | 64446 |
| Longest exon (bp) | 2.17E+04 | 21680 |
| Longest five_prime_utr (bp) | 26237 | 7502 |

| | | |
|---|---|---|
| Longest start_codon (bp) | 3 | 3 |
| Longest stop_codon (bp) | 3 | 3 |
| Longest three_prime_utr (bp) | 34989 | 21469 |
| Longest cds piece (bp) | 18318 | 18318 |
| Longest five_prime_utr piece (bp) | 20564 | 7502 |
| Longest three_prime_utr piece (bp) | 21469 | 21469 |
| Longest intron into cds part (bp) | 477863 | 477863 |
| Longest intron into exon part (bp) | 477863 | 477863 |
| Longest intron into five_prime_utr part (bp) | 452131 | 223885 |
| Longest intron into three_prime_utr part (b | 337929 | 118710 |
| Shortest gene (bp) | 120 | 120 |
| Shortest transcript (bp) | 120 | 120 |
| Shortest cds piece (bp) | 3 | 3 |
| Shortest five_prime_utr piece (bp) | 1 | 1 |
| Shortest three_prime_utr piece (bp) | 1 | 1 |
| Shortest intron into cds part (bp) | 20 | 20 |
| Shortest intron into exon part (bp) | 20 | 20 |
| Shortest intron into five_prime_utr part (bp | 20 | 23 |
| Shortest intron into three_prime_utr part (b | 20 | 20 |
| **BUSCO Completeness (arthropoda)** | | |
| | C:97.2%[S:9.7%,D:87.5%],F:0.8% | C:91.6%[S:87.6%,D:4.0%],F:1.0%,M |
| Complete BUSCOs (C) | 984 | 928 |
| Complete and single-copy BUSCOs (S) | 98 | 887 |
| Complete and duplicated BUSCOs (D) | 886 | 41 |
| Fragmented BUSCOs (F) | 8 | 10 |
| Missing BUSCOs (M) | 21 | 75 |
| Total BUSCO groups searched | 1013 | 1013 |

Γ.

l:7.4%,n:1013

**Table S6.** EnTAP functional annotation statistics for the longest isoforms using 70/70 coverage.

| **Functional Annotation** | **Longest Isoform (70/70 coverage)** |
|---|---|
| Total Input Sequences | 18996 |
| Similarity Search | |
| Total unique sequences with an alignment | 11247 (59.21% of total input sequences) |
| Total unique sequences without an alignment | 7749 (40.79% of total input sequences) |
| Top 10 alignments by species | |
| | 1)centruroides sculpturatus: 7286(64.78%) |
| | 2)limulus polyphemus: 736(6.54%) |
| | 3)stegodyphus dumicola: 613(5.45%) |
| | 4)parasteatoda tepidariorum: 467(4.15%) |
| | 5)ixodes scapularis: 167(1.48%) |
| | 6)dermacentor silvarum: 91(0.81%) |
| | 7)rhipicephalus sanguineus: 90(0.80%) |
| | 8)rhipicephalus microplus: 78(0.69%) |
| | 9)cryptotermes secundus: 26(0.23%) |
| | 10)zootermopsis nevadensis: 25(0.22%) |
| Gene Families | |
| Total unique sequences with family assignment | 14120 (74.33% of total input sequences) |
| Total unique sequences without family assignment | 4876 (25.67% of total input sequences) |
| Total unique sequences with at least one GO term | 12563 (66.13% of total input sequences) |
| Total unique sequences with at least one pathway (KEGG) assignment | 4421 (23.27% of total input sequences) |
| Totals | |
| Total retained sequences (after filtering and/or frame selection) | 18996 |
| Total unique sequences annotated (similarity search alignments only) | 306 (1.61% of total retained) |
| Total unique sequences annotated (gene family assignment only) | 3179 (16.74% of total retained) |
| Total unique sequences annotated (gene family and/or similarity search) | 14426 (75.94% of total retained) |
| Total unique sequences unannotated (gene family and/or similarity search) | 4570 (24.06% of total retained) |

**Table S7.** Existing scorpion genome quality statistics

BUSCO

| Species | BUSCO | Single Copy % | Duplicated % | Fragmented % | Missing % | N50 | contigs | assembly_size |
|---|---|---|---|---|---|---|---|---|
| *M. martensii* | 44.70% | 42.7 | 2 | 38.4 | 16.9 | 45790 | 54037 | 9.14E+08 |
| *C. vittatus* | 97.20% | 92.7 | 4.5 | 1.5 | 1.3 | 2358559 | 2067 | 7.61E+08 |
| *C. sculpturatus* | 94.20% | 89.8 | 4.4 | 3.8 | 2 | 537465 | 8337 | 9.25E+08 |
| *A. mauritanicus* | 68.30% | 54.1 | 14.2 | 23.7 | 8 | 4313 | 460271 | 1.08E+09 |
| *H. arizonensis* | 98.60% | 95.2 | 3.4 | 1 | 0.4 | 2.80E+08 | 9 | 2.23E+09 |

| genome_size | auN |
|---|---|
| 880M | 57847 |
| 880M | 2889993 |
| 1.32 G | 647173.2 |
| 1.1 G | 9074.4 |
| 2.33 G | 2.57E+08 |
